# A Pair of Cadmium-exposed Zebrafish Affect Boldness and Landmark use in the Un-exposed Majority

**DOI:** 10.1101/2023.11.09.566440

**Authors:** Delia S. Shelton, Piyumika S. Suriyampola, Zoe M. Dinges, Stephen P. Glaholt, Joseph R. Shaw, Emília P. Martins

## Abstract

Some individuals have a disproportionate effect on group responses. These individuals may possess distinct attributes that differentiate them from others. These characteristics may include susceptibility to contaminant exposure such as cadmium, a potent trace metal present in water and food. Here, we tested whether a pair of cadmium-exposed individuals could exert an impact on the behavior of the unexposed majority. We used behavioral assessments to characterize the extent of the effects of the cadmium-exposed pair on group boldness, cohesion, activity and responses to landmarks. We found that groups with a pair of cadmium-exposed fish approached and remained closer to novel stimuli and landmarks than did groups with pairs of fish treated with uncontaminated water (control). Shoals with cadmium and water treated fish exhibited similar levels of cohesion and activity. The results suggest that fish acutely exposed to environmentally-relevant cadmium concentrations can have profound effects on the un-exposed majority.

## Main Text

Contaminants are rising in the environment with untold impact on wildlife. Wildlife are exposed to contaminants through anthropogenic forces such as industrialization and sprawling urbanization leading to increased toxic effects on ecosystem health. Some contaminants (e.g., metals, PFAS) persist in the environment for decades [1,2]. Consequently, chemicals have been detected in the carcasses of wide variety of animals [2–4]. Traditionally, toxicology has focused on the lethal effects but has increasingly moved to understanding sublethal effects [5]. These sublethal effects are largely individually-measured morphological and behavioral endpoints, however [5,6]. Many animals spend part of their lives in groups – it is estimated that 75% of fish species school or shoal during their ontogeny [7], 50% of bird species form feeding flocks [8], and a large percentage of insects [9], mammals, amphibians, bacteria [7,8,10], and to a lesser extent reptiles aggregate [11]. Animals form groups to identify landmarks, forage for food, and investigate novel stimuli. Characterizing the effects of contaminant exposure on group living can enhance environmental risk assessments and better predict population declines and ecological instability.

Group living animals reside in aggregations with diverse individuals. Some of these individuals are more susceptible to disease and pollution, whereas others are not [12,13]. For example, mice with low metallothionein are more susceptible to cadmium toxicity, than counterparts that have higher metallothionein [14,15]. Migratory male loggerhead turtles accumulate more organic contaminants than resident male turtles [16]. When contaminated individuals return to the group, they may indirectly expose other members and affect their behavior. For example, returning bee foragers exposed to contaminated food sources exposed other bees and larvae to the insecticide laden pollen [17], and the colonies with exposed individuals grew more slowly than controls [18]. Therefore, it is important to identify the impact of contaminated individuals on uncontaminated individuals in group tasks.

Some individuals with distinct phenotypes or experiences can disproportionately group responses. In collectives, these individuals can alter group cohesion, foraging, navigation [19], and social networks [20,21]. These influential individuals may have a greater disease resistance [22], higher dominance rank [19], or more information [23]. There is evidence that individuals in poor health and expressing deficits may also affect the group [24,25]. Some individuals maybe impaired by infection, toxins or drugs, which then affects the group’s behavior. For example, stickleback pairs with an infected individual were slower, less cohesive and more uncoordinated than pairs of uninfected fish [26]. Zebrafish shoals with a single individual exposed to alcohol swam faster and showed an altered leader-follower relationship in comparison to control shoals [27]. Groups may also respond by moving these animals to periphery of the group, expelling them, or maintaining close associations. For example, infected ants move to the periphery of the social network [28], wasps remove the carcasses of dead and dying wasp from the colony [29], and male finches prefer to preferentially feed near diseased conspecifics [30]. Here, we ask if a pair of individuals impaired by contamination can also affect the group’s response.

One notoriously persistent contaminant is cadmium [31]. Cadmium is a trace metal that is heavily utilized in industrial settings, and through runoff it winds up in aquatic habitats [32,33]. Cadmium toxicity can lead to death and morphological deformities and lower concentrations of cadmium can lead to adverse behavioral effects [2,34]. At the individual level, exposure to acute low concentrations of cadmium leads to reduced activity, increased boldness and deficits in anti-predatory behavior [35]. At the group level, there is a paucity of information on the effects of cadmium on behavior, but some studies show that cadmium exposure alters aggression, social networks, dominance hierarchies, response to migratory cues, antipredatory responses, and boldness [4,36–40]. Expanding our understanding to include how contaminants affect group foraging and cognition will provide us with a comprehensive understanding of the adverse effects of contaminants on basic activities that are essential for group living species to persist.

We examined whether a pair of Cd-exposed individuals could alter the groups response to ecological stimuli. Identifying ecological effects of a pair of polluted individuals on the group’s response in a widely used model organism such as the zebrafish (*Danio rerio*) allows us to characterize contaminant effects that are translatable to other species. Zebrafish are native to possess a well-documented behavioral repertoire that is amenable to high throughput screening [41]. Increasing knowledge of their natural history enables us to create ecologically relevant testing arenas [41–45]. To test for an effect of Cd-contaminated fish on group responses to environmental stimuli, we created shoals of familiar fish. From each group, we assigned two fish to either an acute exposure consisting of an environmentally relevant concentration of Cd or water (or control). We measured differences between groups with control and Cd-exposed zebrafish in relation to their response to a novel stimulus, food, landmarks and shoal members. These behavior are ecologically and evolutionarily important and may forecast how contaminated individuals can modify group responses and how these groups may then respond atypically to environmental stimuli [46–49].

## Method

### Subjects

We used Scientific Hatcheries strain of zebrafish managed by Aquatica Biotech (Florida, USA), which is an outbred, wild-type strain. Several recent studies have used this strain of zebrafish [36,44,50–52]. The fish were maintained in 38 L (10 gallon) glass tanks, at 28° C, under 10:14 hour light/dark cycle and fed flake food *ad libitum*. The water quality parameters were similar to those described in Shelton et al., 2023 [36].

### Procedure

We created 36 groups of adult zebrafish with 6 fish per group. The fish became familiar with each other over at least two weeks. We then selected a male and female fish from each group and placed half the pairs in a 1L glass beaker with system water with 0 μg/L Cd (control) and the others with 1μg/L Cd for 17h overnight. This exposure regime has been shown to induce visual impairments in adult zebrafish [36]. We used an environmentally relevant concentration of Cd in this study [2,53,54]. We then depurated the treated fish by washing them twice in beakers of system water. We then returned the pairs to their groups for behavioral assessments. A more detailed description of the exposure and depuration procedure can be found in Shelton et al., 2023 [36].

To test if a pair of cadmium-exposed fish could influence the group’s response to a novel stimulus, landmarks and food, we placed the groups in the testing arena. The testing arena was a 20.8 L (43×23 cm) glass aquarium with shallow water (8 cm depth). To enhance the automated tracking, we increased the contrast between the fish and the background by fitting the tanks with a white plastic floor. Above the center of the floor, we placed a white balloon that could be inflated to a fixed diameter (3 cm); the inflated balloon served as a novel stimulus. To record the behavioral trials, we positioned a webcam (Logitech® c525 HD) for video-recording at 30 frames/s above the test arena.

We recorded the behavior of the group for 5 mins and then presented a novel stimulus after 4 mins lapsed. After the completion of all trials, we immediately began the feeding trial. The feeding trial consisted of sprinkling 500 mg of food into the corner of the arena and assessing the latency for all fish to approach the food. After the feeding trial we then altered the testing arena by adding landmarks (or plants). We placed 4 small plastic plants (12 cm between plants, occupying 0.7% of the total volume and 3.5% surface area) in the tank. The 8 cm in heigh plants (Green Foreground Plastic Aquarium Plants), were painted with fish safe white paint to decrease contrast to the white plastic floor, thereby enhancing tracking. The plants were similar to the vegetation that we observed in the microhabitats of wild zebrafish [41,43,45] and we used with wild zebrafish in the lab [55]. We permitted the fish to acclimate to the testing conditions before assessing their response to the landmarks on the following morning (about 20 h post manipulation and 1 h after the lights came on).

To automatically track the zebrafish from recordings, we used EthoVision XT10 (Noldus Information Technology 2013) software. We used the software to identify the x and y coordinates of individual fish in 2-dimensional space from above every 0.03s (1790 moments/ min). We then calculated several aspects of social behavior from those points. Overall, EthoVision tracked the fish with a high degree of accuracy, however, we did drop approximately 6.2% of the 5370 moments in each trial because EthoVision was unable to detect each fish. The control and experimental treatment were similar in this proportion. We dropped four videos for the novel stimulus (n = 17 Cd, n = 15 control groups) and 1 video in the plant treatment (n = 18 Cd, 17 control groups) due to technical malfunctions. Feeding trials were scored in real time (n = 18 fish/condition).

We quantified several behavioral responses: 1) latency to the novel stimulus, lapse in time between the inflation of the balloon and a fish from the group touching the balloon, 2) nearest neighbor distance, minimum distance between fish, 3) shoal diameter, maximum distance between fish, and 4) activity or total distance moved (cm/3min) was calculated by summing across the six zebrafish in the shoal at each moment, and then averaging over the 5370 moments in the trial, as described in [44]. We estimated the response to landmarks as proximity of the shoal to plants. We first created a heat map of the activity of the shoals, by assessing the percentage of frames the centroid of the fish was over a particular area in the arena. We then measured the shortest distance from the area of highest percentage of the occurrence of fish (red patch) to the nearest plant. For these fish, we also assessed activity (or distanced moved) of the groups in the presence of landmarks as described above. To compare groups with pairs of Cd-exposed and control fish, we used independent sample’s t-test (two-tailed) for latency to approach novel stimulus, nearest-neighbor distance, shoal diameter, activity, distance to landmark, and latency to feed. We applied Welch’s correction when data were not equal in variance. We used non-parametric Mann-Whitney *U* test when data were not normally distributed and thus violated the t-test assumptions. We used R with the “base” package to conduct all statistical analyses [56].

## Results

### Cadmium-exposed Fish Affect Shoal Boldness

We found that the groups with a pair of cadmium-exposed fish had a shorter latency to approach the novel stimulus than the groups with a pair of control zebrafish. The shoals with cadmium-exposed fish (ME = 0.5, SD = 14.42) took 1.5 s less to approach the balloon than the shoals with water-exposed fish (ME = 2.0, SD = 15.13), which led to a significant difference in latency to approach the novel stimulus (*U* = 92, *p* = 0.01; Fig. 1).

**Figure 1.**
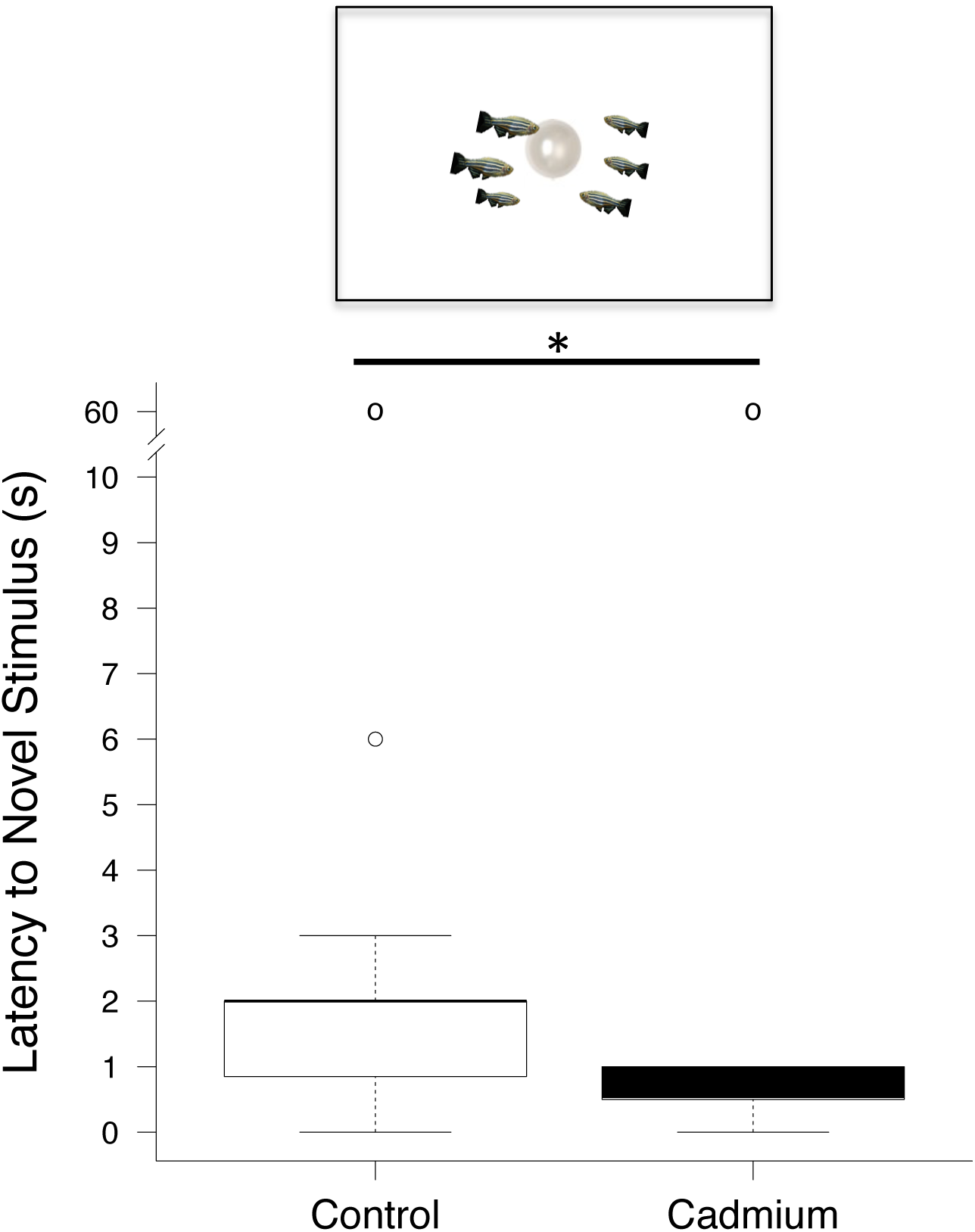
Cadmium pairs altered group boldness. The latency to approach the novel stimulus positioned in the center of the tank. * indicates significant Mann-Whitney *U* comparisons at *p* < 0.05. ° represents outliers. Medians with interquartile ranges (n = 15 for groups with water (or control) treated pairs, and n =17 for groups with cadmium-treated pairs).

### Cadmium-exposed Fish affect Landmark Use, but not Activity of the Un-exposed Majority

To evaluate ecological effects of Cd pairs on group behavior, we assessed group responses to landmarks and food, important cognitive and metabolic correlates [49,57]. We placed groups with Cd- or un-treated pairs in arenas with plants and mapped shoal activity in relation to the landmarks (plants represented as white circles; Fig. 2A). We then assessed the group’s proximity to landmarks or the shortest distance from the shoal to the nearest plant. Groups with Cd-treated pairs remained more than 2 cm or 1.6 times closer to the landmarks than groups with un-treated pairs (Fig. 2B; *U* = 221, *p* < 0.03). In the presence of plants, groups with cadmium-exposed and control fish are equally active (Fig. 2C). Groups with cadmium-exposed pairs and control pairs moved an average of 5362 (SD = 807.5) cm and 5085 (SD = 1095.7) cm in the trial, respectively. This led to a non-significant difference in activity between the experimental and control groups (*t(*28.30) = -0.83, *p* = 0.41).

**Fig. 2.**
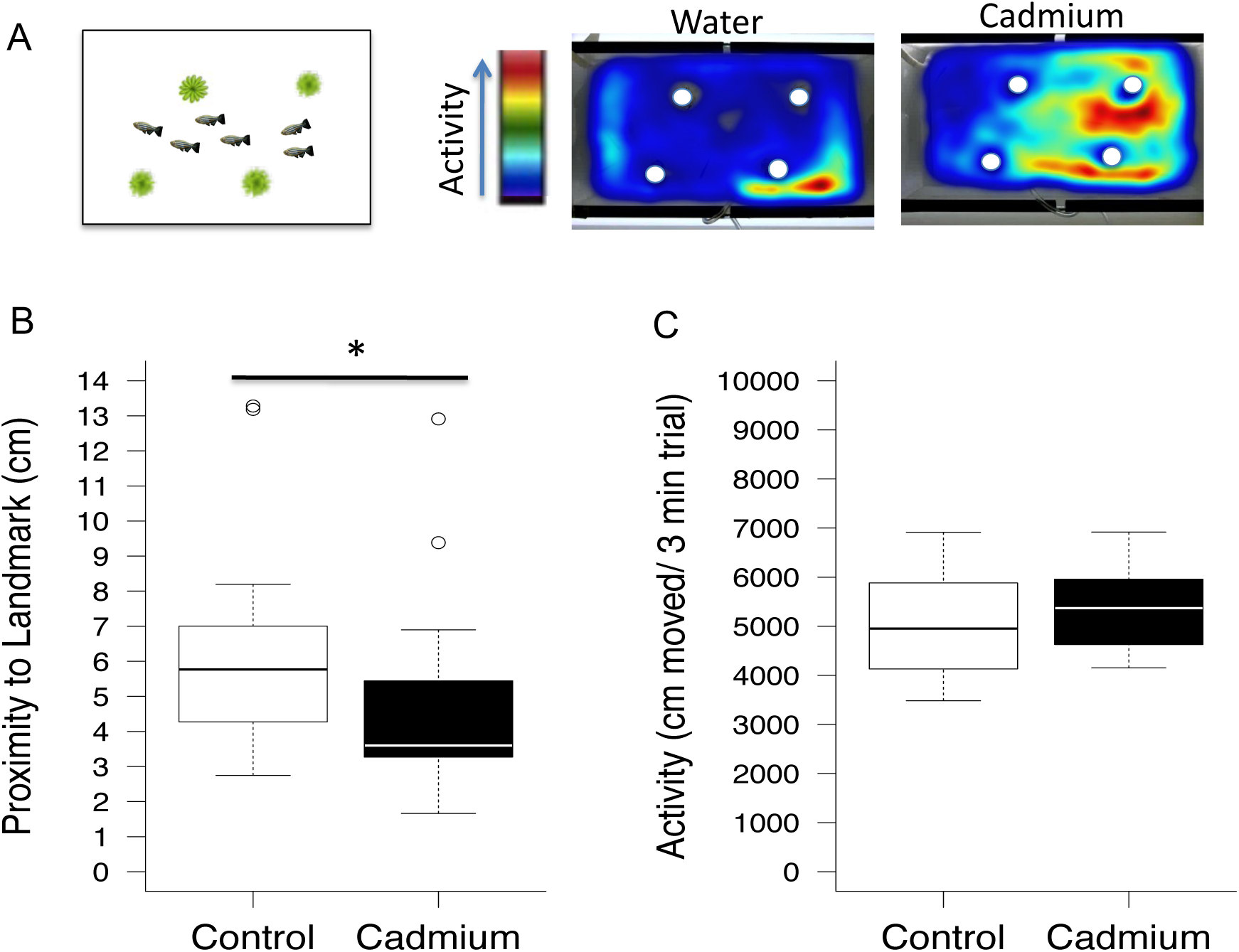
Group responses in the presence of landmarks. (**A**) Representative tank with plant landmarks and heatmaps of group responses to these landmarks. A color gradient shows which activity level corresponds to each color with the lowest level in gray and the highest level in red. (**B**) Groups with cadmium-treated pairs (red patch) are closer to landmarks (white circles) than groups with un-treated pairs. (**C**) Groups with control and cadmium-treated pairs had similar levels of activity (*U* = 123, *p* = 0.33). * indicates significant Mann-Whitney *U* comparisons at *p* < 0.05. ° represents outliers. Medians with interquartile ranges (n = 17 or 18 for each treatment).

### Groups with cadmium and control pairs of fish are equally cohesive and efficient at finding food

The groups were equally cohesive (Fig. 2D, 2E). Groups with cadmium pairs had 7.7 cm NND (SD = 2.60) and a 20.9 (SD = 4.05) cm shoal diameter, and groups with control pairs had 7.6 cm NND (SD = 3.27) and 21.6 (SD = 5.59) cm shoal diameter. These similar individual and group level measures of cohesion led to nonsignificant differences in NND (*U* = 187, *p* = 0.27) and shoal diameter (*t*(29.07) = - 0.43, *p* = 0.67). Groups with Cd pairs displayed a longer apparent latency to feed than those with control pairs, but the difference was not significant (*U* = 163.5, *p* = 0.97; Fig. 3).

**Figure 3.**
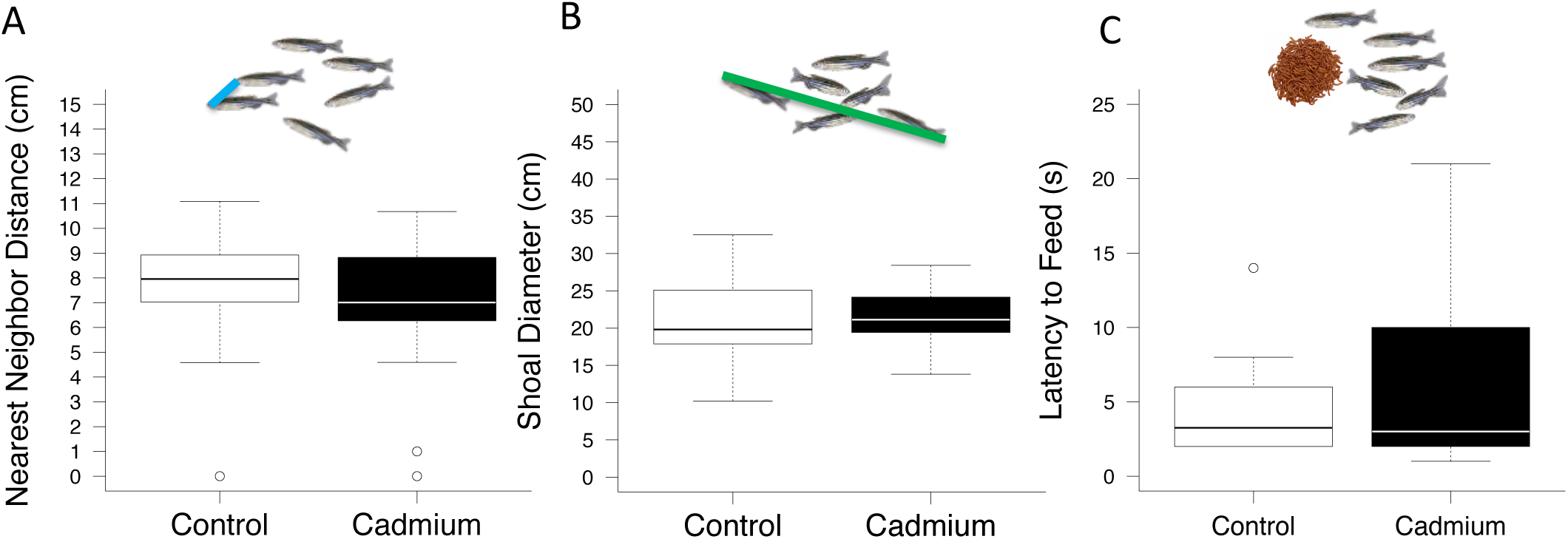
Group cohesion and feeding responses. **A**) Groups had similar nearest neighbor distances (*U* = 187, *p* = 0.27). (**B**). Groups had similar shoal diameters (*t*(29.07) = - 0.43, *p* = 0.67). C) Groups had a similar latency to feed on flake food (*U* = 163.5, *p* = 0.97). Medians with interquartile ranges (n = 17 or 18 for each treatment). ° represents outliers.

## Discussion

Groups of zebrafish with a pair of Cd-treated fish were bolder or approached the unfamiliar object more promptly than did groups with a pair of control fish. Pairs of Cd-treated fish influenced the group’s responses to landmarks, but not their activity, shoal cohesion or foraging behavior. The maintenance of cohesive shoals for groups with a pair of Cd-treated fish suggests that there is benefit for shoaling with deficient individuals. The larger group size may allow the group to compensate so that they can continue to forage typically, but may have adverse effects when encountering novel stimuli in the environment. In the context of anthropogenic change, this finding is intriguing because it suggests that the presence of the contaminated individuals may buffer against some aspects of environmental change, but may also be a “social trap” or may put the group at risk in novel conditions [58,59].

We observed that the shoals with a Cd-exposed pair and control groups were equally cohesive. As the present Cd-exposure regime induces visual deficits [22], it is interesting that the groups with impaired fish continue to shoal, because fish with visuals deficits do not shoal as well as intact fish [60]. It is possible that although the present concentration of Cd disrupts vision [36], it does not impair vision sufficiently enough to lead to shoaling deficiencies, or the individuals are relying on other sensory modalities to shoal. The small size of the arena might have allowed the fish to be sufficiently close, so that at this near distance higher visual acuity was unnecessary. Dose dependent effects of Cd-exposure on visual behavior have been shown in larval zebrafish [61,62] and sensory modality shifts have been observed in adult zebrafish and other fishes exposed to contaminants [52,63].

The maintenance of the deficient Cd-exposed individuals in the group maybe a way to reduce risk and enhance foraging efficiency, as individuals in larger groups have reduced individual predation risk [64] and find food faster [65]. More recent studies have shown that group-size dependent responses are affected by contamination events. Fish in turbid waters have a longer latency to forage and reduce cohesion [66] due to lower visual acuity. This behavioral change is possibly an adaptive response to changes in risk perception. We found that the shoals with Cd-exposed and water-treated fish found food equally as fast, which suggest that the group size was sufficient to allow the groups with Cd-exposed visually impaired fish to overcome a handicap as result of shoaling with these deficient fish. Some suggest that individual abilities cannot compensate for the benefits of grouping [67], and there is an optimal group size [23,68], whereas other suggest that the group composition also influences the beneficial or adverse effects of group living [24]. Future studies should identify in which contexts group size and group composition positively and negatively affect the group.

Animals may form groups of diverse composition to enhance foraging success [69], and reduce predation risk [70]. The mixed composition maybe maintained because some of the individuals are more dominant [19], have experience [71], or are more socially central in the network [21], which then provides advantages to the group. For example, bold shoals and mixed shoals of shy and bold guppies approached a novel feeder faster than shoals completely composed of shy individuals [69]. Groups with experienced Cd-exposed fish are closer to the novel stimulus than groups with experienced water-treated fish [36]. Here, we show that shoals with in-experienced Cd-exposed fish are also closer to the novel stimulus. This suggests that Cd-exposure alone may determine the response of the group to novel stimuli. Previously, we showed that composition was more important for determining boldness [36]. The un-exposed fish may maintain these Cd-exposed individuals in the group because their impairments further dilute the risk of the un-exposed fish [36]. The visual impairment of Cd-exposed fish may lead them to be initial targets during a predation event, and thus embolden the un-exposed fish to engage in riskier behavior or moving closer to novel stimuli. Other animal groups maintain compromised individuals in the group. For example, lame animals in ungulate herds are moved to the periphery where they have an increased rate of encountering predators [72], and male finches prefer to feed near diseased conspecifics [30]. The maintenance of these deficient individuals in the group due to the dilution of risk maybe an evolutionary trap, as the initial benefits of the Cd-treated fish may affect the group adversely in other contexts [58,59,73]. Alternatively, the maintenance of the deficient individuals in the group maybe an evolutionarily stable strategy that enhances the survival of the species [74,75]. Group size is a predictor of foraging success [65] and here we also observed that group size, not composition, influenced foraging behavior. Others have observed that group composition influences foraging success [69], but these individuals varied along a bold-shy continuum not in health. Thus, the effects of composition on foraging may depend on the health or contaminant exposure status of the individuals. Some suggest that the advantages of group living are group size dependent, whereas others suggest that the group composition also influences the benefits of grouping [69,70,76]. Future studies should determine if there is a ratio or a number of contaminated individuals that disrupts the ability of the group to maintain species typical responses.

Groups with Cd-exposed fish were closer to landmarks and novel objects than groups with water-treated fish. The need to be closer to the landmarks and novel objects may be because the fish cannot see as well, therefore the have to come closer to the objects. Other studies have shown that when individual vision is compromised due to pollutants that the fish reduce activity and remain closer to refuge due to increased perception of risk [52,66]. In the present study, there is support for an indirect effect of cadmium exposure where the presence of the Cd-exposed individuals generally increased the exploratory behavior of the fish by diluting the risk. Other studies show that pollutants including Cd increase exploratory behavior. Zebrafish experiencing an identical exposure and grouping paradigm as in the present study were also closer to a different type of novel object, moving Eppendorf tube, and were more socially investigative [36]. Fish exposed to psychiatric drugs had a shorter latency to enter a novel arena or were bolder than control fish [77]. Future studies should determine if the indirect effects of Cd-exposed individuals on the group’s response is due to aberrant behavior induced by the pollutant, a more general effect of having deficient individuals in the group or the effect is occurring through another mechanism of action.

It is notable that the concentration of Cd used in the study (1 µg/L Cd exposure in 17 h) is lower the US EPA specific maximum contaminant level for acute Cd exposures to aquatic life specified in 2001 of 2.0 µg/L Cd exposure in 24 h. It is difficult to compare the present exposure paradigm to the 2016 US EPA criterion of 1.8 µg/L Cd in 1 h for acute exposures and 0.72 µg/L Cd in 4 days because of the large differences in exposure duration [78]. We corroborate other findings that concentrations of Cd that are lower than the 2001 US EPA criterions for aquatic exposures can alter the behavioral response of directly and indirectly exposed fish [36,61,62,79]. Other studies reported that fish exposed to 1 µg/L Cd for 24 h showed altered swimming patterns and increased oxidative stress without significant Cd accumulation [80]. Given that the biological half-life of Cd can span decades [31,81], future studies should determine the long-term or trans-generational consequences of low Cd exposures on indirectly and directly exposed organisms. The present experimental setup and use of automated tracking software makes the study of pollutant-induced personality more amendable to higher-throughput screening. This automation will potentially allow environmental regulatory bodies and others to include behavioral endpoints that span directly and indirectly exposed individuals to better protect wildlife. Future studies should also examine the long-term or fitness consequences of acute and chronic Cd exposure on indirectly exposed fish, as acute and chronic exposures can have differing effects. The inclusion of behavioral endpoints and group responses is important when setting criterions, because behavior is a sensitive indicator of physiological changes, is the first target of evolutionary selective forces, and many animals reside in groups.

## Conclusions

Our study suggests that a pair of Cd-exposed individuals can profoundly affect group responses, which adds support to area of growing research interest [47]. Here, we extend the effect of acute Cd exposures to include indirect effects on the group’s response to ecological features and novel environmental stimuli. Understanding the impact of Cd on behavior is an important endpoint because behavior is a predictor of physiological and fitness consequences that can be potentially rescued with appropriate interventions. Our study emphasizes the importance of using group responses as a notable end point because many animals reside in groups during their life, and disruptions to group behavior can have adverse consequences for other life history events.

## Ethics

We conducted all procedures in accordance with the Institutional Animal Care and Use Committee guidelines at Indiana University under protocols 12-042 and 15-017-9.

## Data accessibility

The data presented in this manuscript can be found on DryRad at https://doi.org/10.5061/dryad.gxd2547rn.

## Authors’ contributions

DSS, PSS and EPM designed the experiment. SG and JS helped with cadmium exposure, DSS and PSS ran the experiments, PSS assisted with behavioral analysis, and DSS wrote the manuscript with substantial input from EPM and feedback on earlier versions from PSS and ZMD.

## Acknowledgments

We thank Anuj Khemka, Delawrence Sykes, Dolores Shelton, Ploypenmas Boyd, Jeffrey R. Kelly, Myra Bower, Halima Amro, David Noakes, Jesualdo Fuentes-G, Alison Ossip-Drahos, Stephanie Campos, Jay Goldberg, Jaime Zúñiga-Vega, Mike Simonich and Monserrat Suárez Rodríguez for thoughtful discussions and helpful comments on earlier versions of this article.

